# Architectural principles for Hfq/Crc-mediated regulation of gene expression

**DOI:** 10.1101/464024

**Authors:** Xue-Yuan Pei, Tom Dendooven, Elisabeth Sonnleitner, Shaoxia Chen, Udo Bläsi, Ben F. Luisi

## Abstract

The global regulator Hfq facilitates the action of regulatory RNAs in post-transcription gene regulation in many Gram-negative bacteria. *In Pseudomonas aeruginosa*, Hfq, in conjunction with the catabolite repression protein Crc, was shown to form a complex that directly inhibits translation of target transcripts during carbon catabolite repression. Here, we describe and validate high-resolution cryo-EM structures of the cooperative assembly of Hfq and Crc bound to a translation initiation site. The core assembly is formed through interactions of two cognate RNAs, two Hfq hexamers and a Crc pair. Additional Crc protomers can be recruited to form higher-order assemblies with demonstrated *in vivo* activity. The structures indicate a distinctive RNA conformation and a pattern of repeating motifs that confer regulatory function. This study not only reveals for the first time how Hfq cooperates with a partner protein to regulate translation but also provides a novel structural basis to explain how an RNA code can guide global regulators to interact cooperatively and regulate many different RNA targets.

## INTRODUCTION

The control of gene expression in many bacteria is fine-tuned through intricate post-transcriptional networks mediated by the action of small regulatory RNAs (Wagner and Romby, 2015). Many of these regulatory molecules require the RNA chaperone Hfq, which protects the RNA against ribonucleases and facilitates their base-pairing interactions with cognate RNA targets (Vogel and Luisi, 2011). Hfq is a member of the Lsm/Sm family and shares with that group an ancient structural core that oligomerizes to form toroidal architectures exposing several RNA-binding surfaces. Crystallographic and biophysical data show that RNA recognition is mediated by distinct interactions with distal, proximal and rim faces (Schumacher et al., 2002; Link et al., 2009; Sauer et al., 2012; Panja et al., 2013), as well as revealing the role of the natively unstructured C-terminal tail in autoregulating RNA binding activities (Santiago-Frangos et al., 2016; 2017).

In the opportunistic pathogen *Pseudomonas aeruginosa*, Hfq acts as a pleiotropic regulator of metabolism (Sonnleitner and Bläsi, 2014), virulence (Sonnleitner et al., 2003; Fernandez et al., 2016; Pusic et al., 2016), quorum sensing (Sonnleitner et al., 2006; Yang et al., 2015) and stress responses (Lu et al., 2016). Many of these roles are likely facilitated through partner molecules, and numerous putative protein interactors of *P. aeruginosa* Hfq have been identified with functions in transcription, translation and mRNA decay (Van den Bossche et al., 2014). In the case of Hfq from *Escherichia coli,* the functionally important partners include RNA polymerase, ribosomal protein S1 (Sukhodolets and Garges, 2003), the endoribonuclease RNase E (Ikeda et al., 2011), polyA-polymerase, and the exoribonuclease polynucleotide phosphorylase (Mohanty et al., 2004; Bandyra et al., 2016). Most likely, these complexes are RNA mediated and affect the colocalisation of the transcriptional, translational and RNA decay machineries (Worrall et al., 2008; Resch et al., 2010; Večerek et al., 2010).

One *P. aeruginosa* protein that was found to co-purify with tagged Hfq is the <u>C</u>atabolite <u>r</u>epression <u>c</u>ontrol protein, Crc (Van den Bossche et al., 2014; Moreno et al., 2015; Sonnleitner et al., 2018). Crc is involved in carbon catabolite repression (CCR) in *Pseudomonas*, a process that channels metabolism to use preferred carbon sources (such as succinate) until they are exhausted, whereupon alternative sources are used (Rojo, 2010). Crc strengthens binding of A-rich target transcripts to the distal side of Hfq (Sonnleitner et al., 2018). In this way, translational repression arises through binding of both Hfq and Crc to A-rich ribosome-binding sequences of mRNAs sensitive to catabolite repression, such as *amiE* mRNA, encoding aliphatic amidase (Sonnleitner et al., 2009; Sonnleitner and Bläsi, 2014). The translationally repressed mRNA, e.g. *amiE*, is then subjected to degradation, which might trigger disassembly of the Hfq/Crc/RNA complex (Sonnleitner and Bläsi, 2014). The non-coding RNA CrcZ, which increases in levels when the preferred carbon source is exhausted, was shown to sequester Hfq, and thereby to counteract Hfq/Crc mediated translational repression of mRNAs related to catabolism (Sonnleitner et al., 2009; Sonnleitner and Bläsi, 2014). CrcZ expression is under control of the alternative sigma factor RpoN and the two-component system CbrA/B (Sonnleitner et al., 2009). Although the signal responsible for CbrA/B activation remains unknown, it is thought to be related to the cellular energy status (Valentini et al., 2014).

In addition to carbon catabolite repression, Hfq and Crc link key metabolic and virulence processes in *Pseudomonas* species. The two proteins affect biofilm formation, motility (O’Toole et al., 2000, Huang et al., 2012, Zhang et al., 2012; Pusic et al., 2016), biosynthesis of the virulence factor pyocyanin (Sonnleitner et al., 2003; Huang et al., 2012), and they have been shown to affect antibiotic susceptibility (Linares et al., 2010; Heitzinger, 2016). Recent ChiP-seq studies indicate that Hfq and Crc have an even broader regulatory impact in *Pseudomonas*. It was shown that these regulators can work in concert to bind many nascent transcripts co-translationally, uncovering a large number of regulatory targets (Kambara et al., 2018).

To gain insight into how *P. aeruginosa* Hfq cooperates with Crc in translational repression of mRNAs, we determined the structure of the complex they form on the Hfq binding motif of the CCR-controlled *amiE* mRNA using cryo-electron microscopy (cryoEM). Our analyses revealed that the components form higher order assemblies and explain for the first time how a recurring structural motif can support the association of Hfq and RNA into cooperative ribonucleoprotein complexes that have key regulatory roles. We observe that the interactions supporting the quaternary structure are required for *in vivo* translational regulation. These findings expand the paradigm for *in vivo* action of Hfq through cooperation with the Crc helper protein and RNA to form effector assemblies.

## RESULTS

### An ensemble of Hfq/Crc/amiE_6ARN_RNA assemblies

For cryo-EM structural studies of the Hfq/Crc/RNA complex, purified recombinant Hfq and Crc proteins were mixed with an 18 nucleotide Hfq binding motif from the translation initiation region of the CCR-controlled *amiE* mRNA, which encodes aliphatic amidase (hereafter, *amiE*_6ARN_). This binding motif comprises of 6 repeats of an A-R-N pattern preferred by the distal face of Hfq. The purified sample of Hfq/Crc/*amiE*_6ARN_, after mild chemical crosslinking, yielded well defined single particles on graphene oxide in thin, vitreous ice. Analysis of the reference free 2D class averages and subsequent 3D classification indicated three principal types of complexes corresponding to different stoichiometries of Hfq (hexamer):Crc:*amiE*_6ARN_ with compositions 2:2:2, 2:3:2 and 2:4:2 (Figure 1). These higher order assemblies are in agreement with recently observed SEC-MALS and mass spectrometry results which excluded a simple 1:1:1 assembly (Sonnleitner et al., 2018). The maps for the complexes are estimated to be 3.12 Å, 3.27 Å and 3.27 Å in resolution, respectively, based on gold-standard Fourier shell correlations (Figure 1 – figure supplement 1). The distribution of the complexes corresponds to roughly 40%, 33% and 26% for the 2:2:2, 2:3:2 and 2:4:2 complexes (Figure 1). The individual crystal structures of Hfq and Crc dock well into the cryoEM densities (Figure 1 and Figure 1 – figure supplement 2), and aside from side chain rotations there are few other structural changes of the components (Figure 1 – figure supplement 2). CryoEM analyses of samples that had not been treated by crosslinking show that the quaternary structure was not affected by the treatment (Table S1).

**Figure 1.**
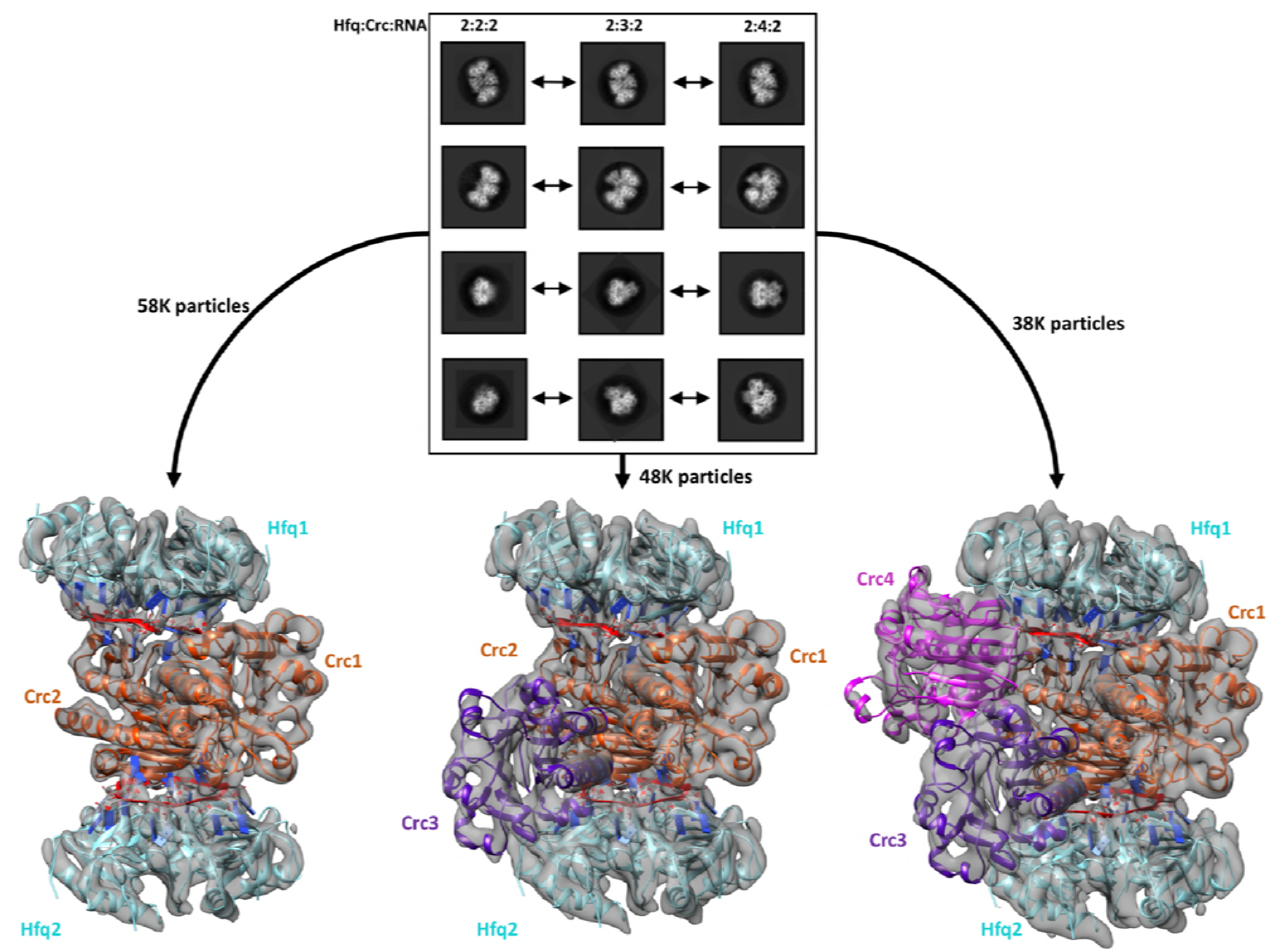
Reference free 2D classification and 3D classification of Hfq:Crc:RNA particles. Three main classes of particles were observed after reference-free 2D classification (top), corresponding to Hfq:Crc:*amiE*_6ARN_ stoichiometries of 2:2:2, 2:3:2 and 2:4:2. The *amiE*_6ARN_ species (red) constitute the main interaction interface between Hfq and Crc, together forming the 2:2:2 core complex observable in all three models (bottom). Cyan: Hfq Hexamer, orange purple and pink: Crc monomers, red: *amiE*_6ARN_. All cryoEM maps were low-pass filtered to 6 Å for interpretability and the crystal structures were docked in as rigid bodies. For clarity, only a subset of all the high quality 2D classes are shown in the panel.

In the core complex (2:2:2), the two Hfq hexamers sandwich the RNA and Crc components (Figure 2A). Each Hfq interacts with one *amiE*_6ARN_ RNA and two Crc molecules, forming an assembly with C2 symmetry. The molecular twofold axis passes through the centre of the two Crc molecules, and the same Crc-to-Crc interface is observed in the crystal structure of the isolated Crc dimer (generated through crystallographic symmetry) (Milojevic et al., 2013). As anticipated, the dominating protein/RNA interaction is made by the distal face of Hfq, forming an interface area of ∼2270 Å^2^. The two Crc molecules interact with RNA residues exposed on the surface of Hfq, and both Crc molecules contact the Hfq-rim on the distal side (Figure 2A).

**Figure 2.**
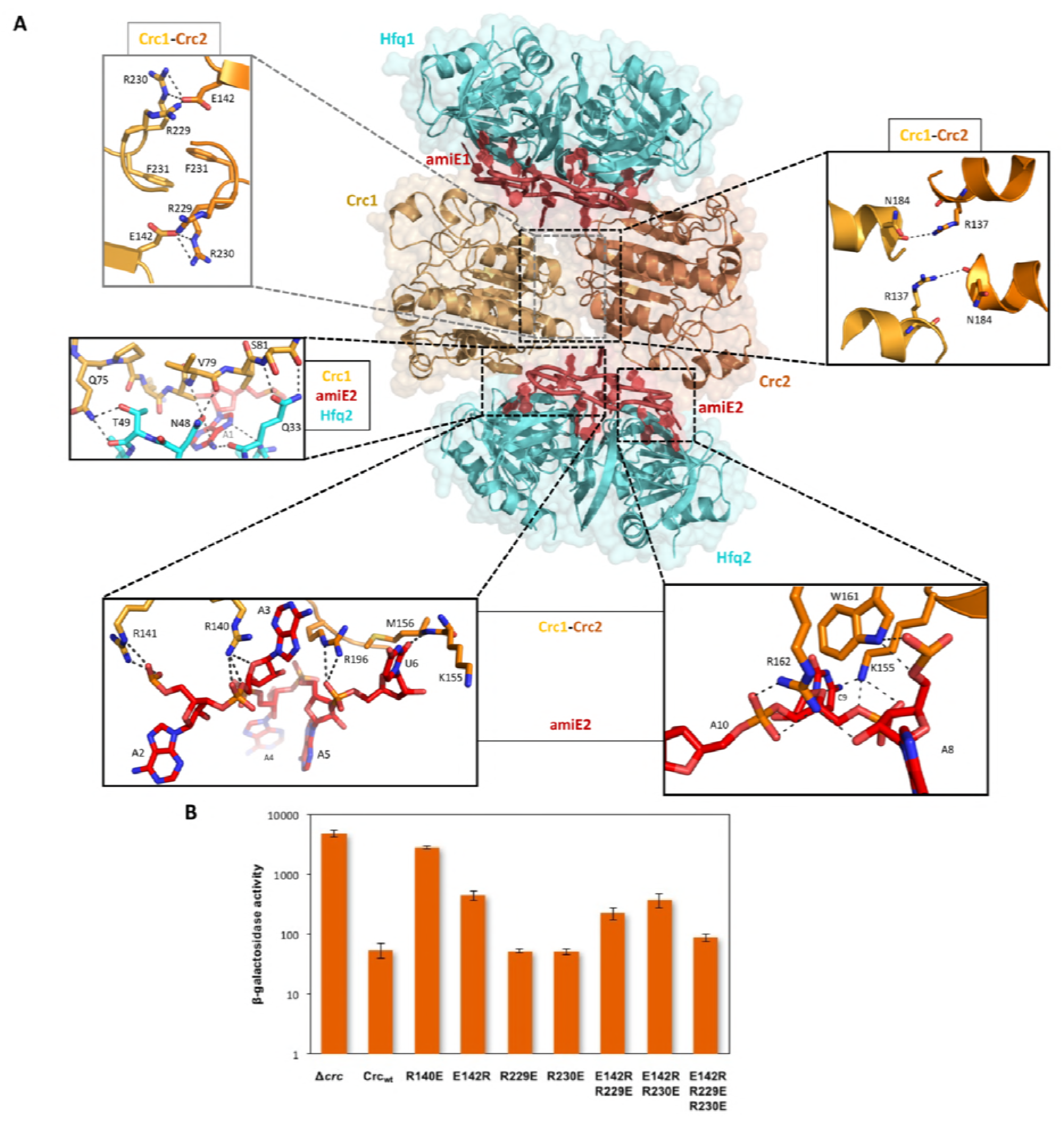
Model of the 2:2:2 Hfq:Crc:RNA complex and validation of interactions. A. Atomic model of the 2:2:2 Hfq:Crc:RNA complex. The view is along the C2 molecular symmetry axis which passes through the homodimeric Crc interface. Hfq hexamers flank the Crc dimer and present the *amiE*_6ARN_ RNA to form two different interfaces with the Crc protomers, which form an anti-parallel dimer. The Crc protomers form strong polar contacts with mainly the backbone phosphate groups and exposed ribose rings (bottom 2 insets). Two small C2 symmetric binding interfaces constitute the Crc dimerisation (top 2 insets). A single short stretch on each Crc monomer binds a Hfq monomer (middle left panel). Dimeric Crc in yellow and orange, *amiE*_6ARN_ RNA in red, Hfq hexamers in cyan. B. Translational repression of an *amiE::lacZ* reporter gene by Crc variants. Strain PAO1*Δcrc*(pME9655) harboring plasmids pME4510 (vector control), pME4510crc_Flag_ (Crc_wt_) or derivatives thereof encoding the respective mutant proteins was grown to an OD_600_ of 2.0 in BSM medium supplemented with 40 mM succinate and 40 mM acetamide. The β-galactosidase values conferred by the translational *amiE*::*lacZ* fusion encoded by plasmid pME9655 in the respective strains are indicated. The results represent data from two independent experiments and are shown as mean and range.

The Crc forms antiparallel dimers in the 2:2:2-complex, so there are two modes of interaction with the *amiE_6ARN_* RNA. Each binding mode is used once at either of the interfaces with Hfq/RNA (Figure 2A). In each Crc monomer, Arg140 η^1^-NH_2_ and Arg141 ε-NH and η^1^-NH_2_ interact with the phosphodiester backbone of *amiE_6ARN_*. Arg140 and Arg196 form a sandwich with the purine-base of the A3 nucleotide at an entry/exit site of *amiE_6ARN_* RNA. The Arg196 ε-NH and η^1^-NH_2_ groups form hydrogen bonds with the U6-*amiE*_6ARN_ backbone and the U6 O2 group forms a hydrogen bond with the Met156 amide. In the second mode of interaction, Lys155 ζ-NH_2_ makes a hydrogen bond with the OP_2_-group of C9 and the ribose hydroxyl group. Additional hydrogen bonds are formed between Trp161 ε^1^-NH and Arg162 η^1/2^-NH_2_ and the phosphate backbone of *amiE_6ARN_* (Figure 2A). These highly organised interactions illustrate how the bases of *amiE*_6ARN_ as presented by Hfq constitute a molecular interface for the RNA-mediated interactions between Hfq and Crc.

The quaternary organisation of the 2:2:2 complex forms a core unit that is also present in the 2:3:2 and 2:4:2 complexes. In that common core, the interaction of the Crc with the RNA leaves approximately half of the accessible surface of the nucleic acid exposed. For the 2:3:2 and 2:4:2 complexes, additional Crc units are recruited through interactions with the exposed portion of the RNA (Figure 3A). As such, the C2 symmetry is broken by the third Crc molecule in the 2:3:2 complex (Figure 1). Interestingly, recruitment of a fourth Crc monomer to the complex restores the C2 symmetry, preserving the symmetry axis from the core complex, but with a conformationally different Crc dimer interface between Crc molecules 3 and 4 (Figure 3A). The two additional Crc monomers have small surface-area contacts with the rest of the complex and are likely to be comparatively mobile, which may account for the stronger variation in resolution for the 2:3:2 and 2:4:2 maps compared to the rather rigid 2:2:2 core assembly (Figure 1-figure supplement 1).

**Figure 3.**
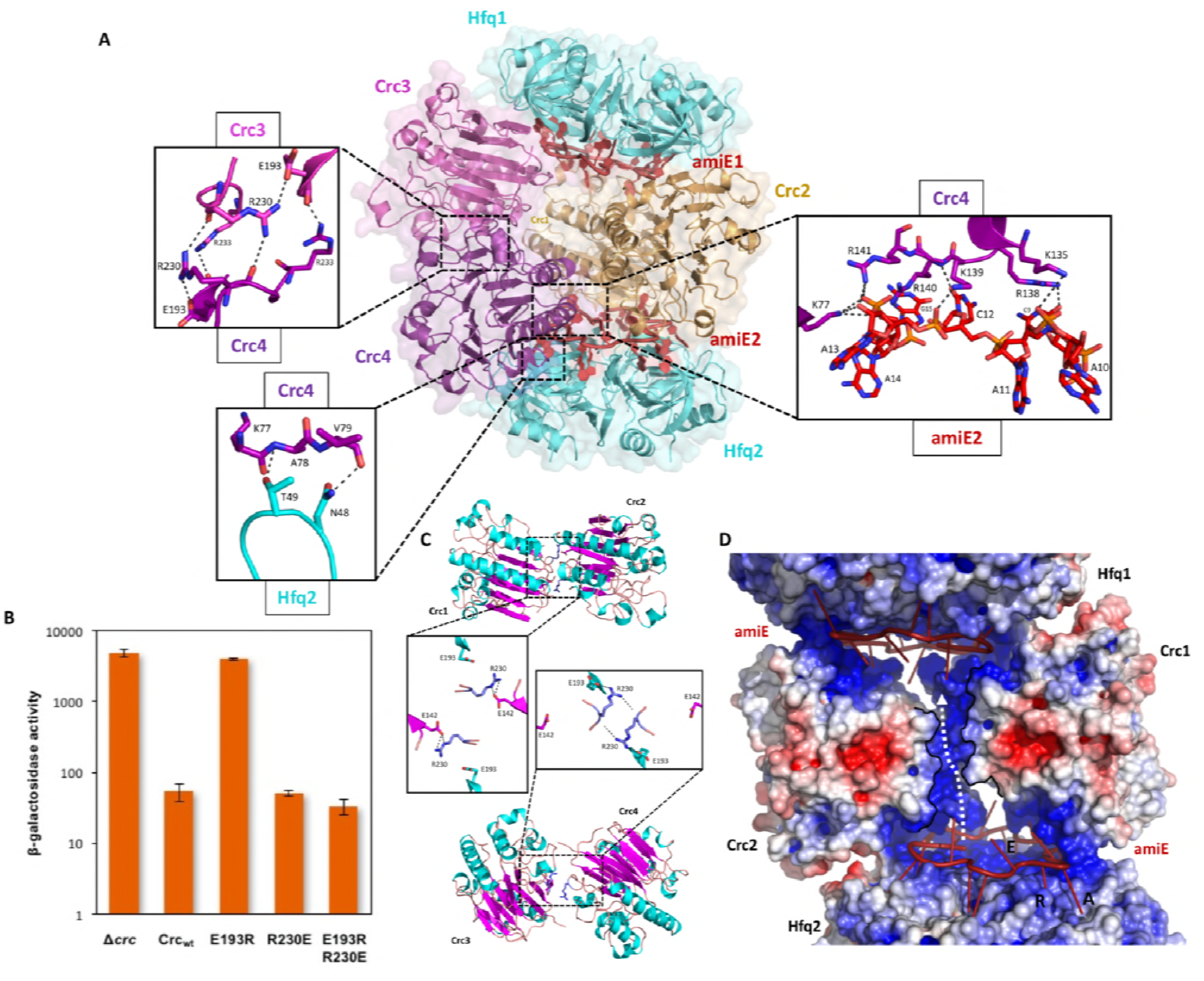
Model of the 2:4:2 Hfq:Crc:RNA complex and validation of interactions. A. Atomic model of the 2:4:2 Hfq:Crc:RNA complex. The insets show additional Hfq-Crc, Crc-Crc and Crc-RNA interactions not present in the 2:2:2 complex. The Crc3-4 dimer is formed by only one interface, with an R230-R230 interaction at the core, which globally overlaps with the dimer interface of the Crc1-Crc2 dimer (top left inset). Only one of two RNA binding patches is presented to *amiE_6ARN_* in the Crc3-4 dimer, yet exploited more extensively (right inset). A small interface is formed between Crc3-4 and Hfq. Crc dimer in yellow, *amiE*_6ARN_ RNA in red, Hfq hexamers in cyan, extra Crc dimer in magenta and purple. B. Translational repression of the *amiE::lacZ* reporter gene by Crc variants, as described in Figure 2B. The results represent data from two independent experiments and are shown as mean and range. C. Two dimeric Crc species are observed over the three complexes solved by cryoEM. i: The self-complementary interaction of the 2:2:2 complex. ii: In the 2:4:2 complex, an alternative dimer is formed, showing a twisted dimer interface and more open configuration, with Arg230 serving as a dynamic hinge (bottom). D. An electropositive half-channel runs along the dimer interface of the Crc1-2 dimer, and in the context of the Hfq/Crc/RNA assembly it could potentially serve as a conduit for RNA (dotted white arrow). The A, R, and E sites are annotated.

### Function, origins and validation of subunit cooperativity in the 2:2:2 complex

Hfq binds the *amiE*_6ARN_ RNA avidly with a dissociation constant in the nanomolar range, but in contrast Crc has no intrinsic RNA-binding activity (Milojevic et al., 2013). In the presence of Crc, the off-rate for Hfq on *amiE*_6ARN_ decreases (Sonnleitner et al., 2018), which indicates a cooperation of the components in binding RNA. The complexes revealed here show that Crc forms small contact surfaces to the RNA, to Hfq, and to itself as a homodimer; these small areas work together to give an assembly that is most likely stabilised through chelate cooperativity. Notably, there is a striking absence of any lower order assemblies in the cryo EM micrographs. The 2:2:2 complex is therefore likely to be the minimal complex formed when all components are present and must be constructed in an ‘all or nothing’ manner, somewhat like a binary switch.

The dimer interface of the Crc pair is the largest protein-protein interface in the 2:2:2-complex and has a buried area of 766 Å^2^, which typically corresponds to a moderate intermolecular affinity. The key dimerization interface is maintained by salt bridges between Arg229-Arg230 of one Crc monomer and Glu142 of the second Crc monomer, which is further stabilised by pi-stacking of the Phe231-Phe231 rings at the point of symmetry (Figure 2A). The phenylalanine residues are in turn stabilised by stacking interactions with Trp255 of the same Crc monomer (not shown). Two additional polar contacts are formed between Arg137 and the Asn184 carbonyl group of two pairs of helices in the Crc dimer, forming a smaller secondary interface (Figure 2A).

The observed interactions shown for the 2:2:2 Hfq:Crc:RNA complex (Figure 2A) are consistent with genetic, biochemical and biophysical data, which revealed intermolecular interactions between Crc protomers, interactions between Crc and RNA as well as a few interactions between Crc and Hfq. These data also showed that formation of the Hfq/Crc/RNA complex requires binding of the RNA on the distal face of Hfq (Sonnleitner et al., 2018). To verify selected interactions between the Crc protomers and Crc and RNA, we explored the effects of mutations on translational repression of an *amiE::lacZ* reporter gene by Hfq and Crc (Sonnleitner et al., 2018), and on the capacity of Hfq and Crc to co-immunoprecipitate in the presence of *amiE*_6ARN_ RNA. First, we asked whether R140 (Crc_R140_) is required for the interaction of the protein with the RNA (Figure 2A, bottom left inset). As shown in Figure 2B, the Crc_R140E_ mutant was deficient in repression of the *amiE::lacZ* reporter gene, similarly as observed in the *crc* deletion strain. Moreover, Crc_R140E_ did not co-immunoprecipitate with Hfq in the presence of *amiE*_6ARN_ RNA (Figure 2 – figure supplement 1), strongly indicating that the interaction between Crc_R140_ and RNA is pivotal for Hfq:Crc:RNA complex formation.

Next, we focused on the possible role of the salt bridges between the E142 and R229/R230 ‘triangle’ (Figure 2A, top left inset) for the Crc-Crc interaction. The single mutant proteins Crc_R229E_ and Crc_R230E_ did not affect translational repression of *amiE::lacZ*, whereas the function of the Crc_E142R_ variant was diminished (Figure 2B), indicating that E142 can form salt bridges with either R229 or R230. The de-repression of *amiE:lacZ* observed with the Crc_E142R_ variant was partially compensated by the double mutant proteins Crc_E142R, R229E_ and Crc_E142R, R230E_. In addition, the Crc_E142R_ and Crc_R230E_ variants were impaired in Hfq:Crc:RNA complex formation as shown by the co-immunoprecipitation assay (Figure 2 – figure supplement 1A). Strikingly, the compensatory changes present in the triple mutant protein Crc_E142R, R229E, R230E_ almost fully restored translational repression of the *amiE::lacZ* reporter gene. As the respective Crc variant proteins were produced at comparable levels (Figure 2 – figure supplement 1B), these mutational studies support the *in vivo* role for the interactions of the Crc protomers observed in the cryo-EM models.

### Function, origins and validation of subunit cooperativity in the 2:4:2 complex

The protomer interactions of the 2:2:2 assembly are highly interdependent, and once the core complex is generated it can recruit additional Crc molecules, forming the 2:3:2 and 2:4:2 complexes. In the 2:4:2 complex, a second type of Crc dimer seems to assemble with a smaller buried surface (Figure 3A). Such additional dimers can only form when an intact 2:2:2 core complex is present, as they are not observed in the core complex itself nor in solution or through crystallographic symmetry (Milojevic et al., 2013). Notably, the additional dimer is a more ‘open’ conformation of the crystallographic Crc dimer in the core, which is further supported by normal mode analysis (data not shown). The same key Crc dimer interface is occupied but seems to serve as a dynamic hinge, whereas the secondary, smaller, dimer interface between the Crc helices is absent to allow the new Crc dimer to adopt an ‘open’ conformation. Arg230 is reorganised by Glu193 in the same protomer to self-interact with the corresponding Arg230 in the partner Crc, rather than with Glu142 (Supplementary movie 1). Additional hydrogen bonds are formed between Arg233 and Glu193, whereas Arg229 is no longer part of the dimer interface (Figure 3A). Both Arg230 and Glu193 seem to play pivotal roles in providing the structural freedom to form a dynamic hinge (Figure 3C).

Only the Arg233-Glu193 interaction is unique for the 2:4:2 assembly and was assessed *in vivo*. Strikingly, Crc_E193R_ fully abrogated repression of the *amiE:LacZ* reporter gene (Figure 3B). The model predicts that the deleterious Crc_E193R_ mutation can be compensated by the substitution of Crc_R230E_ to re-establish the interaction. This pair does indeed behave as predicted, further confirming the *in vivo* importance of the 2:4:2 assembly during CCR (Figure 3B). By reorganising the extra Crc molecules 3 and 4 that bid the 2:2:2 core (Figure 3A), the alternative Crc dimer is able to utilise one of two basic patches on its surface when engaging *amiE*_6ARN_ without causing steric hindrance to the already bound crystallographic dimer.

In addition to Crc Arg140 and Arg141, Crc K139 ζ-NH_2_ makes a hydrogen bond with the OP_2_- group of A12, Arg138 η^1^-NH_2_ interacts with the ribose hydroxyl group of C9 and K135 ζ-NH_2_ forms a hydrogen bond with the A11 OP_2_. Finally, the O2 of cytosine C12 engages in a hydrogen bond with the backbone amino group of Arg140. Direct interactions between the reorganised Crc dimer and Hfq are limited to the same Crc β-strand and exposed loop of a sole Hfq monomer, as in the core complex. Due to the open conformation of the alternative Crc dimer, the Hfq Thr49 hydroxyl group now forms a hydrogen bond with the Ala 78 amide group (Figure 3A).

Interestingly, a basic half-channel is formed over the core dimer interface, with additional basic patches spread over the RNA binding surface of the Crc dimer (Figure 3D). Speculatively, longer RNA species could travel though the surface exposed half-channel and interconnect all components of the core complex into a highly organised assembly on this target RNA.

### A specialised and recurring RNA conformation in Hfq-mediated regulation

Link et al. (2009) described the crystal structure of *E. coli* Hfq bound to a polyriboadenylate 18-mer and observed that the RNA encircled the distal face of the Hfq hexamer *via* a repetitive tripartite binding scheme. Each base triplet is partially embedded between adjacent Hfq monomers and is mostly surface exposed, folding into a ‘crown-like’ conformation. We observe striking similarities with the fold of the authentic *amiE*_6ARN_ species on the distal side of the *P. aeruginosa* Hfq hexamer (Figure 4). Notably, the cryoEM maps were calculated without any reference to the Link et al. (2009) structure. The agreement between the co-crystal structure of the homologous complex and the entirely independently derived cryoEM based model is a strong validation of both experimental procedures, X-ray crystallography and cryoEM. A recent study proposed an RNA-RNA stacking interface between two RNA species presented by Hfq, supported by crystal structures and biophysical analysis in solution (Schulz et al., 2017). Although all components necessary for such interaction are present in our reaction mixture, we do not observe such dimeric species by cryo-EM or in solution when Crc is present.

**Figure 4.**
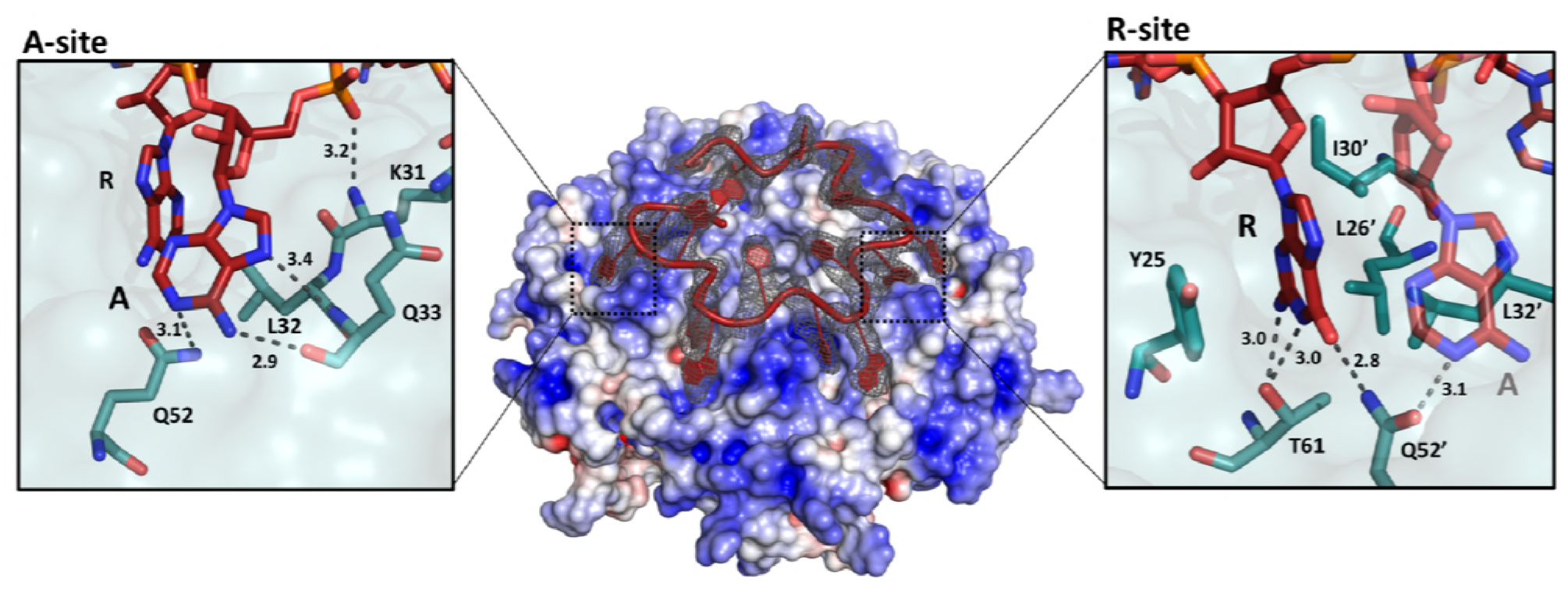
The ‘A-R-N crown’ in the Hfq/*ami*E_6ARN_ RNA complex. 6 RNA triplets are partially embedded in 6 binding pockets on the Hfq distal side, forming a weaving, crown-like pattern. The A and R sites are occupied by adenine and a purine, respectively, whereas the RNA entry/exit site has no discriminatory preferences. Cryo-EM density for *amiE_6ARN_* is depicted as a grey mesh, with the RNA ‘crown’ modelled in red. Positively charged protuberances (blue) guide the RNA to fold into a star-shaped conformation to maximize the surface interaction between the negatively charged RNA backbone, and the positively charged Hfq surface pattern. An atomic model of the A-R-E occupation pattern. Left panel: Adenosine specificity site. Right panel: Purine specificity site. *amiE* nucleotide carbon atoms are depicted in red, Hfq carbon atoms are in green.

Like its *E. coli* homologue, *Pseudomonas* Hfq contains 6 tripartite binding pockets on the distal side, capable of binding a total of 18 nucleotides. Each of the six RNA triplets of the *amiE*_6ARN_ RNA fits into an inter-subunit cleft in Hfq (Figure 4). The specific, star-shaped RNA fold is guided by six positively charged protuberances on the distal face of Hfq, with the phosphate backbone circularly weaving in between these, seemingly to minimise steric hindrance while maximizing surface interactions (Figure 4). As described by Link et al. (2009), each pocket consists of an adenosine specificity site (A), a purine nucleotide specificity site (R), and a presumed RNA entrance/exit site (E) which is non-discriminatory. Hfq thus has a structural preference for (ARN)_n_ RNA stretches on its distal side, where N is any nucleotide. The adenosine specificity (A) sites are organised identically to the corresponding A sites in *E. coli* Hfq, forming hydrogen bonds between the peptide backbone and carboxyl-groups of Gln33 and the N6,7 atoms of the adenosine base, and a polar interaction between Gln52 (N_ε_) and the N1 atom of the adenosine base. The peptide backbone amide of residue Lys31 interacts with the 5’ phosphate group of adenine. Finally, the adenine base is stacked against the side chain of Leu32 (Figure 4). The purine (R) specificity site is defined by two neighbouring monomers, where the side chains from Tyr25 and from Leu26’, Ile30’ and Leu32’ (where the prime denotes residues from a neighbouring subunit) contact the nucleotide aromatic base. In *amiE_6ARN_*, one R-site is populated by a guanine, forming a hydrogen bond between the N_ε_ of Gln52’ and the guanine exocyclic O6 (Figure 4). Just like in the *E. coli* Hfq/polyA_18_ structure (Link et al., 2009), Gln52’ forms a physical link between the A and R sites. Previous structures were obtained from polyA RNA, whereas the structures presented here were solved with the authentic *amiE* Hfq recognition site. Interestingly, Thr61 O_γ_ forms a double hydrogen bond with the N1 and the exocyclic N2 from the guanine base, which was not seen previously (Link et al., 2009) as all R-sites were occupied by adenine residues (Figure 4).

## Discussion

Many functional studies have highlighted how global posttranscriptional regulators cooperate with each other and their RNA targets to control the fate of transcripts with high specificity. A major gap in our current understanding has been the lack of high resolution structural data of these highly coordinated cellular processes. Here we report the first atomic model of Hfq interacting with a translational initiation region (*amiE*_6ARN_) and a partner protein to form a multi-component assembly that mediates translational control (Kambara et al., 2018; Sonnleitner et al., 2018). The RNA is a recurring A-rich fragment of *amiE* that occupies almost entirely the distal surface of Hfq, weaving in between basic, surface exposed islands. There are striking similarities to the structure of the polyA_18_ complex with *E. coli* Hfq reported by Link et al. (2009), whose structure greatly added to the understanding of RNA binding and chaperone mechanisms, and hinted at how the distinct polyA RNA interaction might enable Hfq-mediated regulation. The polyA/Hfq structure revealed rules for recognition of motifs of the type A-R-N, where R is purine and N is any base. The *P. aeruginosa* Hfq interaction with *amiE*_6ARN_ follows the same rules. The A-R-N repeat occurs in many RNAs, and it has been proposed that the exposed bases could mediate RNA to RNA interactions (Schulz et al., 2017). It is also a recurring motif in the nascent transcripts that are associated with Hfq and Crc in *Pseudomonas* (Kambara et al., 2018). We observe that the exposed bases (entry/exit site) and RNA backbone in the Hfq/*amiE*_6ARN_ complex are available for interactions with Crc to form a cooperative assembly that efficiently mediates catabolite repression *in vivo* when the preferred carbon source is available (Figure 5).

**Figure 5.**
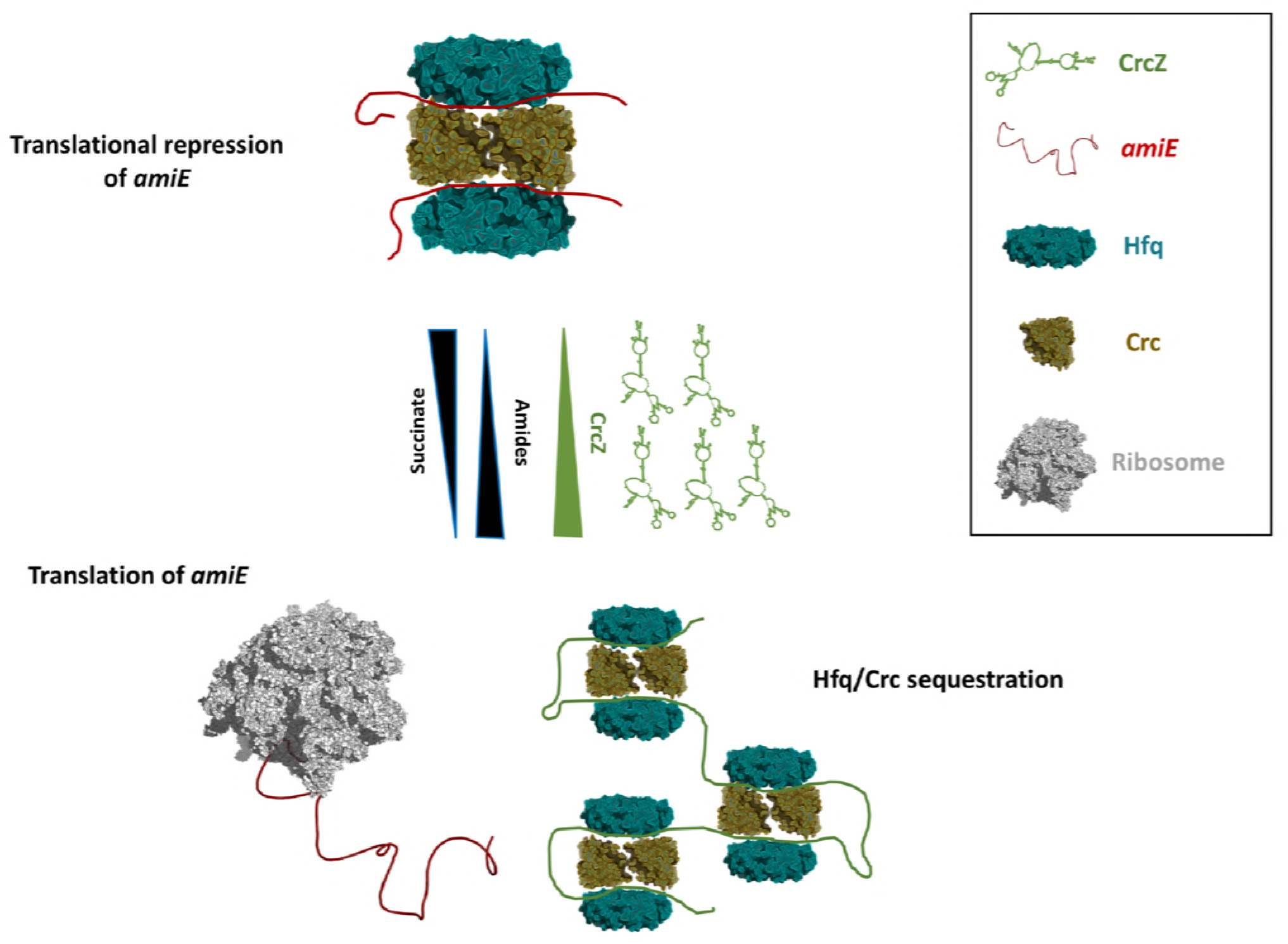
Schematic pathway of the Carbon Catabolite Repression. When the preferred carbon source, succinate, is abundant, cellular CrcZ levels are low and Hfq and Crc occlude the *amiE* ribosome binding site by forming a higher order assembly, rendering CCR active and repressing synthesis of aliphatic amidase (top). Upon depletion of succinate, CrcZ levels increase and sequesters Hfq and Crc from *amiE*, potentially by occupying the multiple ARN patches on CrcZ and forming multicomponent ‘beads on a string’. As such CCR is deactivated, allowing metabolism of a secondary carbon source, *e.g.* amide conversion by AmiE.

Previous studies have shown that both Hfq and Crc are required for tight translational repression of mRNAs, which are subject to carbon catabolite repression (CCR) (Sonnleitner and Bläsi, 2014; Moreno et al., 2015). The presence of Crc did not significantly enhance the affinity of Hfq for *amiE*_6ARN_ RNA (Sonnleitner et al., 2018). However, the simultaneous interactions of Crc with both binding partners resulted in an Hfq/Crc/RNA assembly with increased stability when compared with the Hfq/RNA complex alone (Sonnleitner et al., 2018). In light of our structural studies, the enhancing effect of Crc in Hfq-mediated translational repression of target mRNAs during CCR (Sonnleitner and Bläsi, 2014; Moreno et al., 2015) can be readily explained by the interactions of Crc with both binding partners. It also accounts for the observed decrease in the off-rate on the RNA substrate (Sonnleitner et al., 2018). It is conceivable that full repression is only achieved when *amiE_6ARN_* is masked entirely in the 2:4:2 complex, which is supported by our *in vivo* studies.

The question arises why a higher order assembly such as the 2:2:2 core is formed and not a simpler complex. The structural data indicate that the dimerization of Crc provides the key step for formation of the 2:2:2 complex, because it will pre-organise a copy of the surface that interacts with the Hfq/RNA so that a second Hfq/RNA complex can be recruited. Thus, all components are necessary to form the complex so that there is no formation of lower order ‘sub assemblies’. The structural data are consistent with Crc having no capacity for RNA binding by itself (Milojevic et al., 2013). The Hfq/Crc/RNA complex is thus assembled in a checklist-like manner through numerous small contacting surfaces and when the RNA target is presented by Hfq in a specific, well-defined configuration. In this way, the components interact mutually through chelate cooperative effects. Most likely the 2:2:2 core forms first, then the other Crc components are recruited.

We envisage that the 2:2:2 core and higher order assemblies might interact with other longer RNAs. The higher order assembly could capture two of such mRNA substrates (Figure 5), but chelate effects might instead induce formation of the complex on a single mRNA target. In that scenario, a portion of the mRNA would thread through the central basic half channel as depicted in Figure 3D. Under conditions of catabolite repression regulation, pull-down assays showed that Hfq and Crc form a co-complex in the presence of the 426nt long CrcZ RNA (Moreno et al., 2015; Sonnleitner et al., 2018). In the presence of less preferred carbon sources, the expression levels of CrcZ RNA increase (Sonnleitner et al., 2009) and CrcZ functions as an antagonist in Hfq/Crc mediated translational repression of catabolic genes. The CrcZ RNA has multiple ARN triplets that could be sites for Hfq/Crc interaction (Sonnleitner and Bläsi, 2014) that could sequester multiple Hfq/Crc proteins (Figure 5). Thus, under conditions where CCR is relieved, CrcZ RNA would serve as a sponge for Hfq/Crc to prevent repression of genes encoding proteins required for utilizing the less preferred carbon sources (Figure 5). How the CrcZ RNA is displaced from Hfq/Crc remains unknown. However, the assemblies are likely to be dynamic and the displacement process might resemble that proposed for the step-wise exchange of sRNAs on Hfq (Fender et al., 2010). Recent findings show that the regulatory spectrum of Hfq and Crc is much broader than initially expected. Hfq was found to bind more than 600 nascent transcripts co-transcriptionally often in concert with Crc (Kambara et al., 2018). These findings indicate that Hfq and Crc together regulate gene expression post-transcriptionally beyond just catabolite repression.

Understanding how gene expression is regulated post-transcriptionally in pathogens such as *P. aeruginosa* may provide potential targets for novel drug design. Hfq and Crc are involved in key metabolic and virulence processes in *Pseudomonas* species (O’Toole et al. 2000; Sonnleitner et al., 2003; Sonnleitner et al., 2006; Linares et al., 2010; Huang et al., 2012; Zhang et al. 2012; Zhang et al., 2013; Sonnleitner and Bläsi, 2014; Pusic et al., 2016). Disrupting the interface of the core assembly of the Hfq/Crc complex might be one strategy to counter, among other, metabolic regulation and consequently its downstream processes that impact on virulence during infection. A recent study showed how overproduction of the aliphatic amidase AmiE strongly reduced biofilm formation and almost fully attenuated virulence in, amongst others, a mouse model of acute lung infection (Clamens et al., 2017). Novel drugs that specifically counteract Hfq:Crc:*amiE* assembly formation and prevent repression of AmiE production could induce the phenotype described by Clamens et al (2017). The high resolution structures presented here provide a starting point for novel strategies to interfere with e.g. carbon regulation in a pathogenic bacterium for therapeutic intervention of threatening infections.

## Acknowledgements

The coordinates and cryoEM maps have been deposited in the PDB and the EMBD. BFL, XYP and TD are supported by the Welcome Trust (200873/Z/16/Z). TD is also supported by an AstraZeneca Studentship. U.B. and E.S. are supported by the Austrian Science Fund (FWF) (www.fwf.ac.at/en) [P28711-B22]. We thank our colleagues Jamie Blaza, Dima Chirgadze, Jiri Sponer, Miroslav Kreply, Kasia Bandyra, Steven Hardwick, Sjors Scheres, Joerg Vogel, Armin Resch and Nguyen Thi Bach Hue for advice, helpful discussions and support. For access and help at facilities, we thank Giuseppe Cannon and staff at the MRC-LMB EM Facility and Kasim Sader at Thermo Fisher Scientific Pharma CryoEM Facility, Nanoscience Centre of University of Cambridge.

## MATERIALS AND METHODS

### Protein synthesis, purification and complex formation

*P. aeruginosa* Hfq and Crc were produced in *E. coli* and purified as described by Sonnleitner et al. (2018). The synthetic 18-mer *amiE*_6ARN_ RNA (5’-AAAAAUAACAACAAGAGG-3’) used in these studies consists of six tripartite binding motifs (Sonnleitner and Bläsi, 2014). The Hfq/ Crc/ RNA complex was prepared by first heating the *amiE*_6ARN_ RNA at 95°C for 5 minutes followed by 50°C for 10 minutes and 37°C for 10 minutes. The RNA was then incubated with the Hfq hexamer at a 1:1 molar ratio on ice for 20 minutes to form a binary complex, then an equal molar ratio of Crc was added. The mixture was incubated on ice for 30 minutes prior to fractionation by size exclusion chromatography using a Superdex 200 column equilibrated in running buffer composed of 20 mM HEPES, pH 7.9, 10 mM KCl, 40 mM NaCl, 1 mM MgCl_2_, and 2 mM TCEP (tris(2- carboxyethyl)phosphine). The peak fractions were buffer exchanged into 20 mM HEPES, pH 7.9, 10 mM KCl, 40 mM NaCl, 5 mM MgCl_2_. Samples used for cross-linking were incubated with bis(sulfosuccinimidyl)suberate (BS^3^) at 150 μM for 30 minutes on ice, followed by quenching at 37.5 mM Tris-HCl pH 8.0.

### CryoEM specimen preparation and data acquisition

Graphene oxide grids are prepared as described by Pantelic et al. (2010). Briefly, 2 mg/ml of graphene oxide solution in water (Aldrich) was diluted ten times in water. After removing aggregation by spinning for 30 seconds at 300 rcf, 2 μl of graphene oxide solution was loaded on freshly glow discharged quantifoil Au-grids (R1.2/1.3, 300 mesh). Glow discharge was performed prior to graphene oxide coating at 45 mA for 60 second with an Edward Sputter Coater S150B at 0.2m Bar at 0.75 KV. After the graphene oxide had been adsorbed for 1 minute, the grids were washed 3 times with 20 μl water, then air-dried for 1 hour at room temperature prior to sample application. Specimens for cryoEM analysis were prepared by applying 2 μl of a 0.65 μM solution of the Hfq/Crc/RNA complex to the Quantifoil Au grids freshly coated with graphene oxide. After an adsorption time of 60s, the grids were blotted for 10 seconds at a blot force of 5, then plunge frozen into liquid ethane using a Vitrobot (FEI). Images were recorded on a Krios G2, Falcon III direct electron detector at 300 kV operating in counting mode (Supplementary Table 3).

### Movie processing, single particle analysis, 3D reconstruction and refinement

Whole frame motion correction was performed on movies with motioncorr2 with dose weighting followed by CTF estimation using gctf (Zhang, 2016; Zheng et al., 2017). RELION-2.1 was used for data processing (Scheres, 2012). Final resolution estimates were calculated after the application of a soft binary mask and phase randomisation and determined based on the gold standard FSC=0.143 criterion (Scheres and Chen, 2012; Chen et al., 2013).

For the BS^3^ treated complex, after manually picking 3159 particles and using suitable references for autopicking, 482426 particles were used for early classifications. After three rounds of rejecting particles by 2D classification, 215774 particles were used for initial model generation and 3D classification. An initial model was generated using an SGD algorithm based on a small subset of particles with diverse orientations (Punjani et al., 2017). During 3D classification, three different complexes were resolved after 25 iterations with an angular sampling of 7.5°: 2Hfq:2Crc:2*amiE_6ARN_* (2:2:2), 2Hfq:3Crc:2*amiE_6ARN_* (2:3:2) and 2Hfq:4Crc:2*amiE_6ARN_* (2:4:2). To properly separate, validate and refine the 3 classes, the same 3D classification was rerun with the new 2:3:2 model as reference model, lowpass filtered to 20 Å resolution. C2 symmetry was observed and imposed for the 2:2:2 and 2:4:2 complexes. Each of the classes was then refined to sub-3.5 Å resolution, followed by per-particle frame alignment for movement correction and per-frame damage weighting. The resulting ‘polished’ particles were subjected to a final refinement round with solvent flattening. All reference models were lowpass filtered to 60 Å prior to refinement. The dominant class (2:2:2) had a resolution of 3.12 Å. Local resolution calculations were done with the relion local resolution estimator (Supplementary Figures 1 and 2A, Supplementary Table 1).

Crystal structures for *P. aeruginosa* Crc (PDB code 1U1S) and Hfq (PDB code 4JG3) were manually docked into the EM density map as rigid bodies in Chimera (Pettersen et al., 2004). The RNA 18-mers were manually built into the density using Coot (Emsley et al., 2010). Refmac5 and Phenix real-space refinement with global energy minimization, NCS-restraints, group B-factor and geometry restraints were used to iteratively refine the multi-subunit complexes at high resolution, followed by manual corrections for Ramachandran and geometric outliers in Coot (Supplementary Table 1) (Emsley et al., 2010; Murshudov et al., 2011; Afonine et al., 2012). Model quality was evaluated with Procheck in CCP4 and MolProbity (Williams et al., 2018). *In silico* 2 Å maps were generated from the atomic models and FSC validation against the experimental maps was performed with the EMDB Fourier shell correlation server (EMBL-EBI) (Figure 1 – figure supplement 2 B).

### Bacterial strains and plasmids

The strains, plasmids and oligonucleotides used in this study are listed in Supplementary Tables S2 and S3.

### Construction of plasmids encoding Crc variant proteins for *in vivo* translational repression assay

To test the proficiency of Crc mutant proteins to co-repress translation of a translational *amiE:lacZ* reporter gene, derivatives of plasmid pME4510*crc*_Flag_ (Supplementary Table S2) were constructed by means of Quick change site directed mutagenesis (Agilent Technologies). Plasmid pME4510*crc*_Flag_ was used together with the corresponding mutagenic oligonucleotide pairs (Supplementary Table S3). The parental plasmid templates were digested with *Dpn*I and the mutated nicked circular strands were transformed into *E. coli* XL1-Blue, generating plasmids pME4510crc_(R140E)Flag_, pME4510crc_(E142R)Flag_, pME4510crc_(R229E)Flag_, pME4510crc_(E193R)Flag_, pME4510crc_(R230E)Flag_, pME4510crc_(E142R, R229E)Flag_, pME4510crc_(E193R, R230E)Flag_, pME4510crc_(E142R, R230E)Flag_ and pME4510crc_(E142R, R229E, R230E)Flag_.

### *In vivo* translational repression of an amiE::lacZ reporter gene in the presence of Crc variants

The ability of the Crc mutant proteins to repress translation of an *amiE::lacZ* reporter gene was tested in a PAO1 *crc* deletion strain bearing plasmids encoding the wt protein or the respective protein variants (Supplementary Table S2) as described by Sonnleitner et al. (2018). The β-galactosidase activities were determined as described (Miller, 1972). The β-galactosidase units in the different experiments were derived from two independent experiments.

### Construction of plasmids employed for the production of selected Crc mutant proteins

The R140E, E142R, R230E single aa exchanges in Crc were obtained by using the QuickChange site-directed mutagenesis protocol (Agilent Technologies). The plasmid pETM14lic-His_6_Crc (Supplementary Table S2) was used together with the corresponding mutagenic oligonucleotide pairs (Supplementary Table S3). The entire plasmids were amplified with Pfu DNA polymerase (Thermo Scientific). The parental plasmid templates were digested with *Dpn*I and the mutated nicked circular strands were transformed into *E. coli* XL1-Blue, generating plasmids pETM14lic-His6Crc_R140E_, pETM14lic-His6Crc_E142R_ and pETM14lic-His6Crc_R230E_

### Purification of Crc and Crc variants

The Crc protein and the Crc variants Crc_R140E_, Crc_E142R_ and Crc_R230E_ were purified from *E. coli* strain BL21(DE3) harboring either plasmid pETM14lic-His Crc or the respective derivatives using Ni-affinity chromatography, followed by removal of the His_6_-tag with GST-HRV14-3C ‘‘PreScission’’ protease as described by Milojevic et al. (2013).

### *In vitro* co-IP studies

The co-IP studies in the presence of 40 pmol of Hfq-hexamer, 120 pmol of Crc protein or of the respective Crc mutant proteins and 40 pmol *amiE*_6ARN_ RNA were performed as described (Sonnleitner et al., 2018).

### Western blot analyses

Equal amounts of proteins were separated on 12% SDS-polyacrylamide gels, and then electro-blotted onto a nitrocellulose membrane. The blots were blocked with 5% dry milk in TBS buffer, and probed with rabbit anti-Hfq (Pineda) and rabbit anti-Crc (Pineda) antibodies, respectively. Immuno-detection of ribosomal protein S1 served as a loading control. The antibody-antigen complexes were visualized with alkaline-phosphatase conjugated secondary antibodies (Sigma) using the chromogenic substrates nitro blue tetrazolium chloride (NBT) and 5-Bromo-4-chloro-3-indolyl phosphate (BCIP).

---

**Figure 1 – figure supplement 1.**
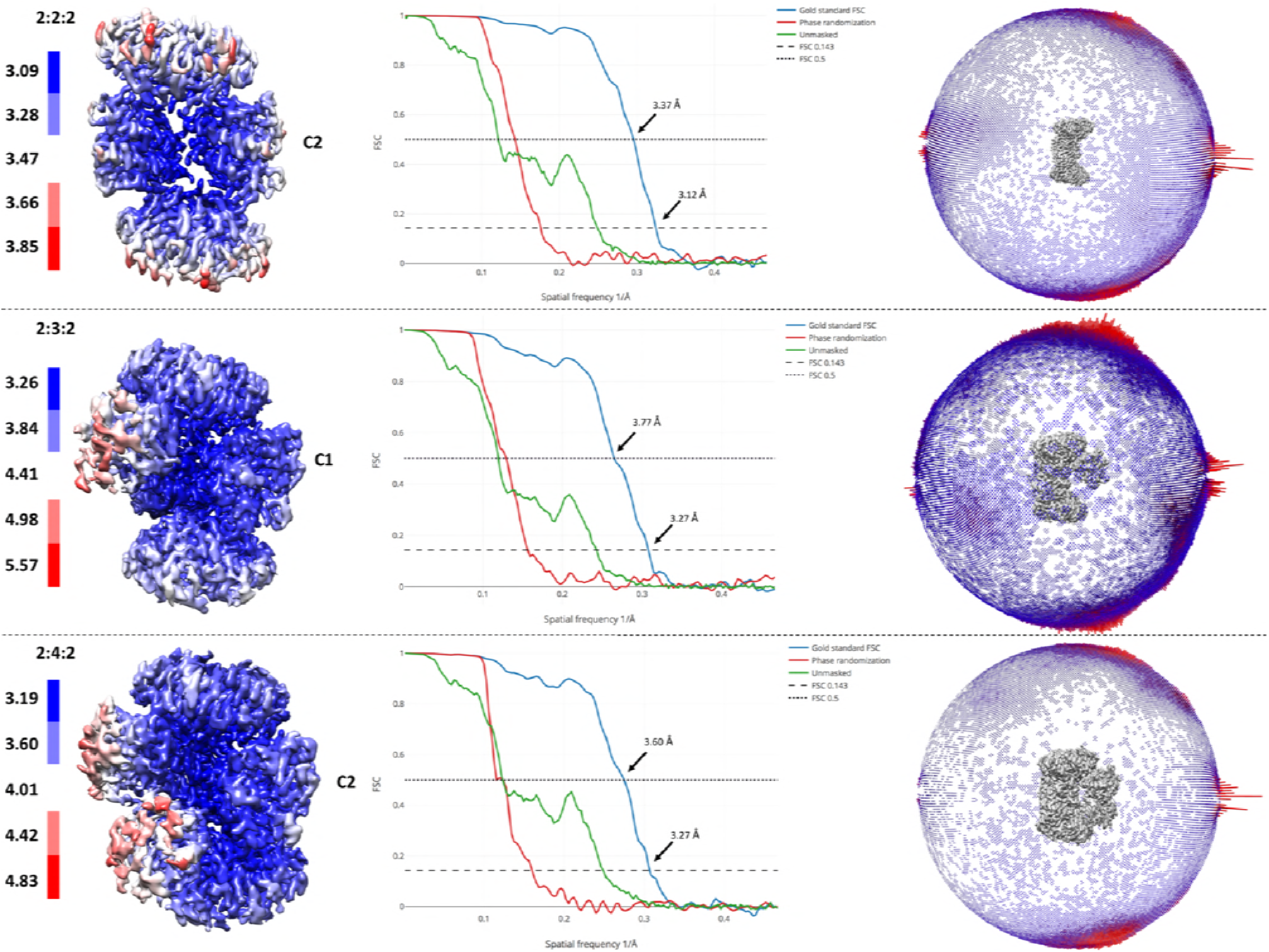
Resolution estimates across the EM maps and Gold standard Fourier shell correlation curves for all three reconstructions. FSC 0.143 and 0.5 are annotated. Angular distributions of the 2D images are presented as a spherical bar plot, with red bars representing more preferred projections.

**Figure 1 – figure supplement 2.**
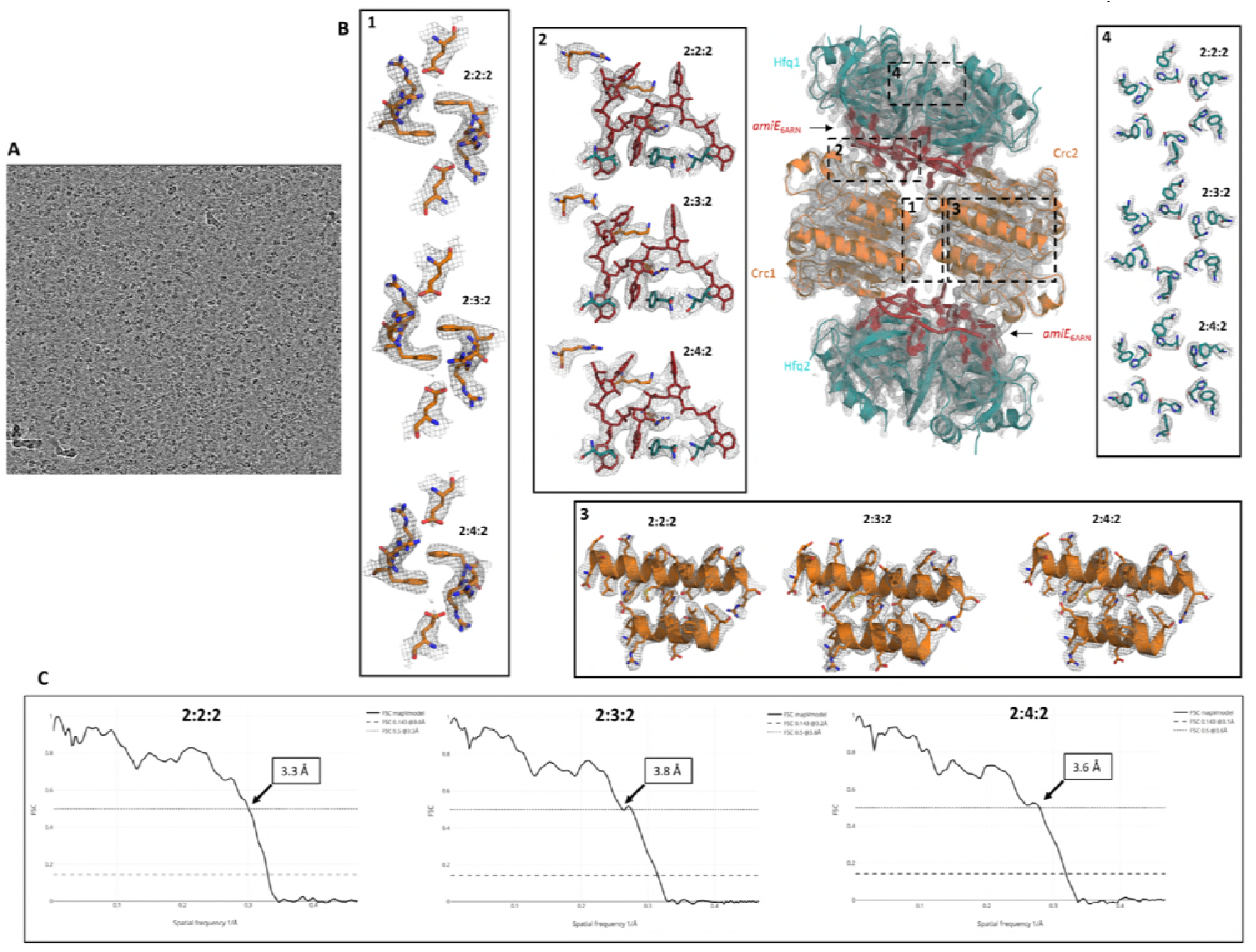
A. Raw micrograph after motion correction at 3 microns under focus. B. High resolution cryo-EM map with refined atomic models for all complexes showing the quality of the EM reconstructions. All maps and models were generated and refined independently of each other and a high-resolution reference structure, showing well defined and highly reproducible densities for all side chains in these signature regions. Even at the periphery the map density is of good quality, maintaining the sixfold symmetry of the Hfq components (inset 4). C. Model versus map Fourier shell correlation (FSC) show a good correlation between the individual atomic models and the experimental cryo-EM maps. FSC 0.5 is annotated on the graph, whereas FSC 0.143 is annotated in the legends.

**Figure 2 – Figure Supplement 1.**
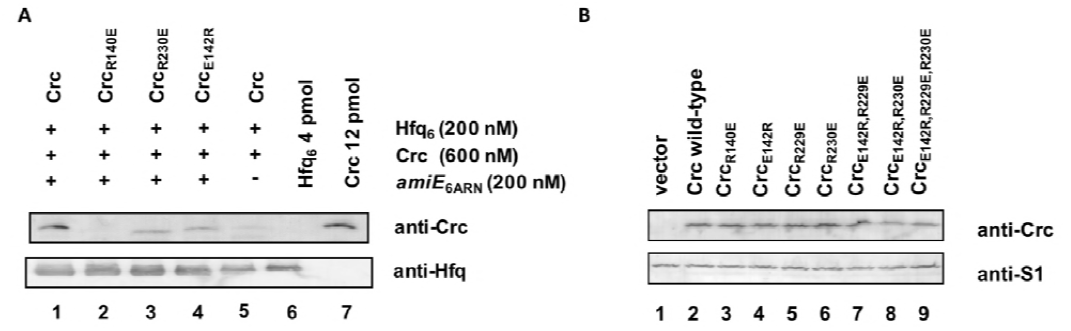
A. *In vitro* association of Hfq and Crc and Crc variants in the presence of RNA. The *in vitro* co-IP experiments were performed with Hfq and Crc and variants in the presence (lanes 1-4) and absence (lane 5) of *amiE*_6ARN_ RNA as indicated on top. Anti-Hfq specific antibodies and magnetic protein G beads were used for co-IP of Crc and Crc variants. The *in vitro* association of Hfq with Crc and variants thereof was visualized by western-blot analysis using anti-Crc or anti-Hfq specific antibodies as indicated at the right. Lane 5, control experiment in the presence of Hfq and Crc but in the absence of RNA. Lanes 6 and 7, 4 pmol Hfq and 12 pmol Crc were loaded, respectively. The western-blot analyses were performed in triplicate. The result from one representative experiment is shown. B. Crc variants are synthesized at comparable levels. Cultures of PAO1Δ*crc*(pME9655,pME4510) (lane 1), PAO1Δ*crc*(pME9655,pME4510crc_Flag_) (lane 2), PAO1Δ*crc*(pME9655, pME4510crc_(R140E)Flag_) (lane 3), PAO1Δ*crc*(pME9655,pME4510crc_(E142R)Flag_) (lane 4), PAO1Δ*crc*(pME9655,pME4510crc_(R229E)Flag_) (lane 5), PAO1Δ*crc*(pME9655,pME4510crc_(R230E)Flag_) (lane 6), PAO1Δ*crc*(pME9655, pME4510crc_(E142R,R229E)Flag_) (lane 7), PAO1Δ*crc*(pME9655, pME4510crc_(E142R,R230E)Flag_) (lane 8) and PAO1Δ*crc*(pME9655, pME4510crc_(E142R,R229E,R230E)Flag_) (lane 9), respectively, were grown to an OD_600_ of 2.0 in BSM medium supplemented with 40 mM succinate and 40 mM acetamide. The protein levels of Crc and Crc variants (top) and of ribosomal protein S1 (loading control) were determined by quantitative western-blot analysis using anti-Crc and anti-S1 antibodies, respectively. The western-blot analyses were performed in triplicate. The result from one representative experiment is shown.

**Supplementary Table S1.** Cryo-EM data collection and refinement statistics for Hfq/Crc/RNA structures

|  |  |  |  |  |  |  |  |
| --- | --- | --- | --- | --- | --- | --- | --- |
| 754 |  |  |  |  |  |  |  |
| 755 | **Supplementary Table S1.** Cryo-EM data collection and refinement statistics for Hfq/Crc/RNA |  |  |  |  |  |  |
| 756 | structures |  |  |  |  |  |  |
| 757 |  | BS<sup>3</sup> crosslinked |  |  | Native |  |  |
| 758 | PDB codes | 6FUS,6FYN,6FXZ |  |  |  |  |  |
| 759 | EMDB codes | 4320, 4326, 4325 |  |  |  |  |  |
| 760 |  |  |  |  |  |  |  |
| 761 | **Data collection** |  |  |  |  |  |  |
| 762 | EM equipment | Titan Krios G2, FEI |  |  | Titan Krios G2, FEI |  |  |
| 763 | Voltage (kV) | 300 |  |  | 300 |  |  |
| 764 | Detector | Falcon III |  |  | Falcon III |  |  |
| 765 | Pixel size (Å) | 1.07 |  |  | 1.09 |  |  |
| 766 | Electron dose (e-/Å<sup>2</sup>/fraction) | 0.37 |  |  | 0.40 |  |  |
| 767 | Defocus range, step (µm) | -1.25 to -3, δ=0.25 |  |  | -1.25 to -3, δ=0.25 |  |  |
| 768 |  |  |  |  |  |  |  |
| 769 | **Reconstruction** |  |  |  |  |  |  |
| 770 | Software | RELION v 2.1 |  |  | RELION v2.1 |  |  |
| 771 | Complex (Hfq:Crc:RNA) | 2:2:2 | 2:3:2 | 2:4:2 | 2:2:2 | 2:3:2 | 2:4:2 |
| 772 | Molecular mass (kDa) | 192 | 222 | 252 | 192 | 222 | 252 |
| 773 | Number of particles used | 57,660 | 48,144 | 38,18 | 25,408 | 18,898 | 14,900 |
| 774 | Angular accuracies (°) | 0.82 | 1.13 | 1.26 | 1.65 | 1.71 | 1.58 |
| 775 | Offsets (pixels) | 0.361 | 0.491 | 0.527 | 0.652 | 0.681 | 0.657 |
| 776 | Symmetry. | C2 | C1 | C2 | C2 | C1 | C2 |
| 777 | Final resolution (Å) | 3.12 | 3.27 | 3.27 | 4.42 | 4.5 | 4.43 |
| 778 | Map-sharpening B factor (Å<sup>2</sup>) | -109 | -125 | -115 | -189 | -172 | -173 |
| 779 | Non-hydrogen atoms | 11466 | 13647 | 15770 | 11466 | 13647 | 15770 |
| 780 | Protein residues | 1292 | 1458 | 1820 | 1292 | 1558 | 1820 |
| 781 | RNA bases | 18 | 18 | 18 | 18 | 18 | 18 |
| 782 |  |  |  |  |  |  |  |
| 783 | **Refinement** |  |  |  |  |  |  |
| 784 | Software | PhenixRSRef |  |  |  |  |  |
| 785 | Model-to-Map CorrelationCoef. | 0.84 | 0.81 | 0.81 |  |  |  |
| 786 |  |  |  |  |  |  |  |
| 787 |  |  |  |  |  |  |  |
| 788 | **Model Validation** |  |  |  |  |  |  |
| 789 | MolProbity score | 1.73 | 1.83 | 1.72 |  |  |  |
| 790 | EMRinger | 3.8 | 3.4 | 3.2 |  |  |  |
| 791 | All-atom clash score | 5.5 | 7.2 | 6.1 |  |  |  |
| 792 |  |  |  |  |  |  |  |
| 793 | **Ramachandran statistics (%)** |  |  |  |  |  |  |
| 794 | Favored (overall) | 93.32 | 93.3 | 94.4 |  |  |  |
| 795 | Allowed (overall) | 6.52 | 6.7 | 5.6 |  |  |  |
| 796 | Outlier (overall) | 0.16 | 0.0 | 0.0 |  |  |  |
| 797 | **R.m.s. deviations** |  |  |  |  |  |  |
| 798 | Bond length (Å ) | 0.008 | 0.023 | 0.007 |  |  |  |
| 799 | Bond angle (°) | 0.89 | 1.09 | 0.86 |  |  |  |
| 800 | **Validation (RNA)** |  |  |  |  |  |  |
| 801 | Correct sugar puckers (%) | 89 | 89 | 89 |  |  |  |
| 802 | Good backbone conformation (%) | 39 | 39 | 39 |  |  |  |
| 803 |  |  |  |  |  |  |  |

**Supplementary Table S2.**
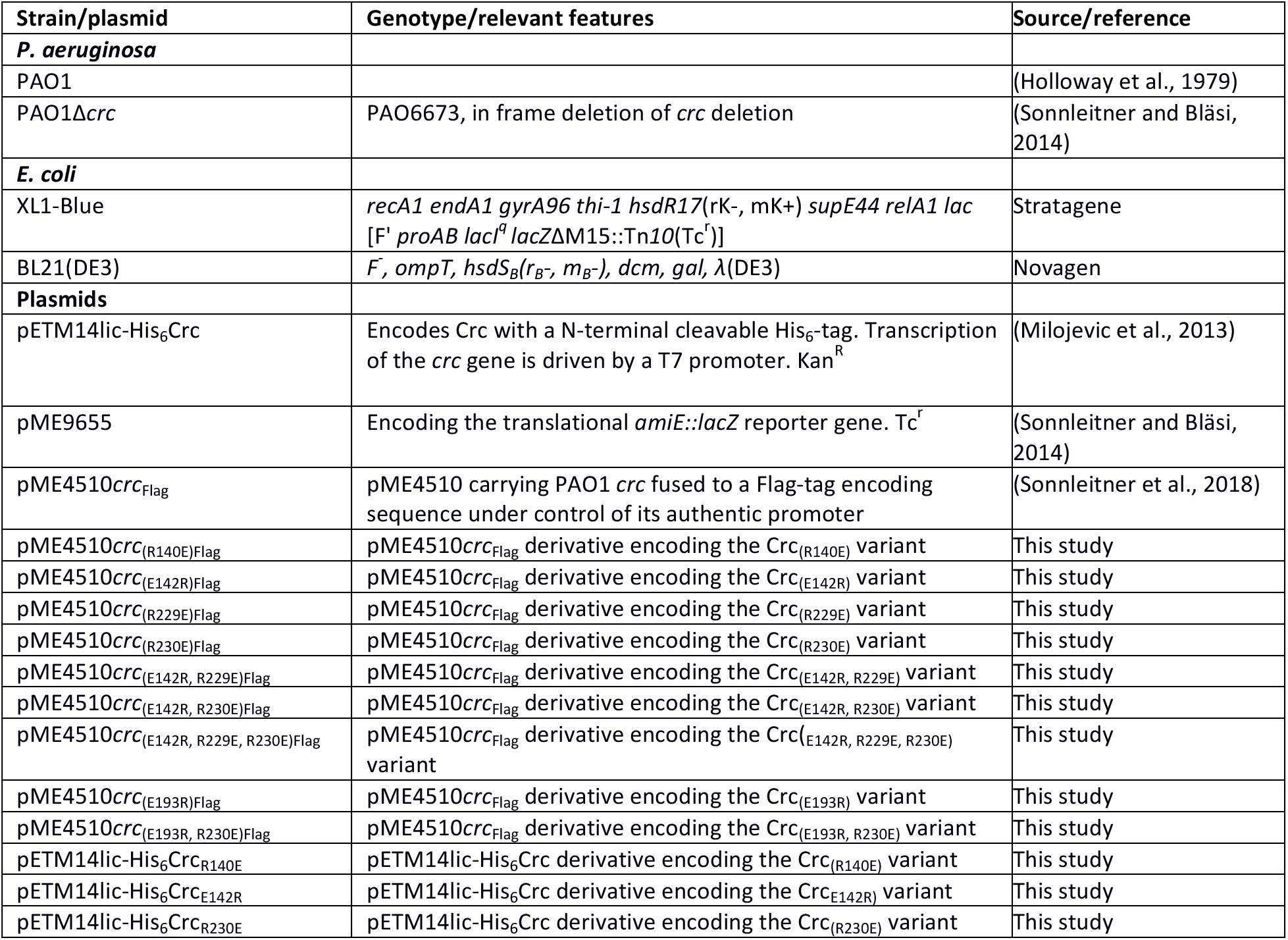
Strains and plasmids used in this study

**Supplementary Table S3.**
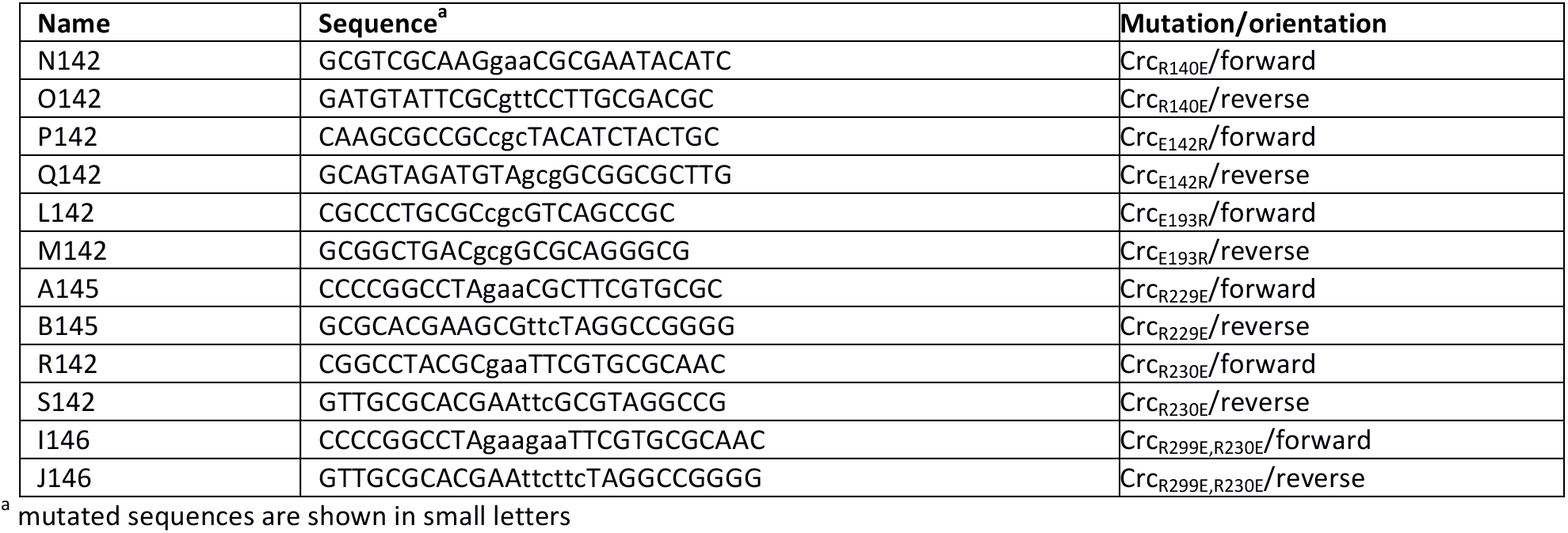
Oligonucleotides used in this study.

